# Identification of insecticidal double-stranded RNAs targeting proteasome subunits in the hop flea beetle *Psylliodes attenuata* using dsRIP

**DOI:** 10.64898/2026.10.23.759960

**Authors:** Doga Cedden, Gözde Güney, Paul Plathner, Michael Rostás, Stefan Scholten, Christina Krönauer

**Affiliations:** Institute for Insect Biotechnology, Justus Liebig University, Heinrich-Buff-Ring 26–32, 35392 Giessen, Germany; Branch Bioresources, Fraunhofer Institute for Molecular Biology and Applied Ecology IME, Ohlebergsweg 12, 35392 Giessen, Germany; Department of Evolutionary Developmental Genetics, Göttingen Center for Molecular Biosciences, University of Göttingen, Johann-Friedrich-Blumenbach Institute, Justus-von-Liebig-Weg 11, 37077 Göttingen, Germany; Department of Animal Physiology, Zoological Institute and Museum, University of Greifswald, Felix-Hausdorff-Straße 1, 17489 Greifswald, Germany; Division of Agricultural Entomology, Department of Crop Sciences, Georg August University of Göttingen, Grisebachstraße 6, 37077 Göttingen, Germany; Division of Crop Plant Genetics, Department of Crop Sciences, University of Göttingen, Von-Siebold-Straße 8, 37075 Göttingen, Germany; Hop Research Center Huell, Institute for Crop Science and Plant Breeding, Bavarian State Research Center for Agriculture, Hüll 5 1/3, 85283 Wolnzach, Germany

**Keywords:** *Psylliodes attenuata*, RNA interference, double-stranded RNA, proteasome, siRNA, dsRIP

## Abstract

The hop flea beetle, *Psylliodes attenuata*, is a major pest of hop (*Humulus lupulus*), an important crop for the brewing industry. Restrictions on chemical insecticides and insecticide resistance create a need for alternative control strategies. As an alternative, RNA interference (RNAi)-based pest control uses double-stranded RNA (dsRNA) to silence essential pest genes, resulting in insecticidal effects. In this study, we aimed to establish *P. attenuata* as an RNAi-based pest control target. We generated a transcriptome assembly using a public RNA sequencing dataset and used the dsRIP web platform to identify essential gene targets and design effective dsRNAs. Three dsRNAs targeting proteasome subunits were tested in feeding bioassays by applying them to hop leaf disks. The treatments caused significant mortality of up to 64% and feeding inhibition of up to 45%. Sequencing of small RNAs associated with the RNA-induced silencing complex identified predominantly 21-nucleotide small interfering RNAs produced from the orally delivered dsRNAs, supporting activation of the RNAi pathway. Additional RNA sequencing of adults enabled assembly of a higher-quality transcriptome with 89% BUSCO completeness for future RNAi studies. Overall, the feeding bioassays and molecular resources establish *P. attenuata* as a promising target for RNAi-based pest control and demonstrate the usefulness of dsRIP for RNAi research in insects previously unstudied at the molecular level.

**Graphical abstract:** 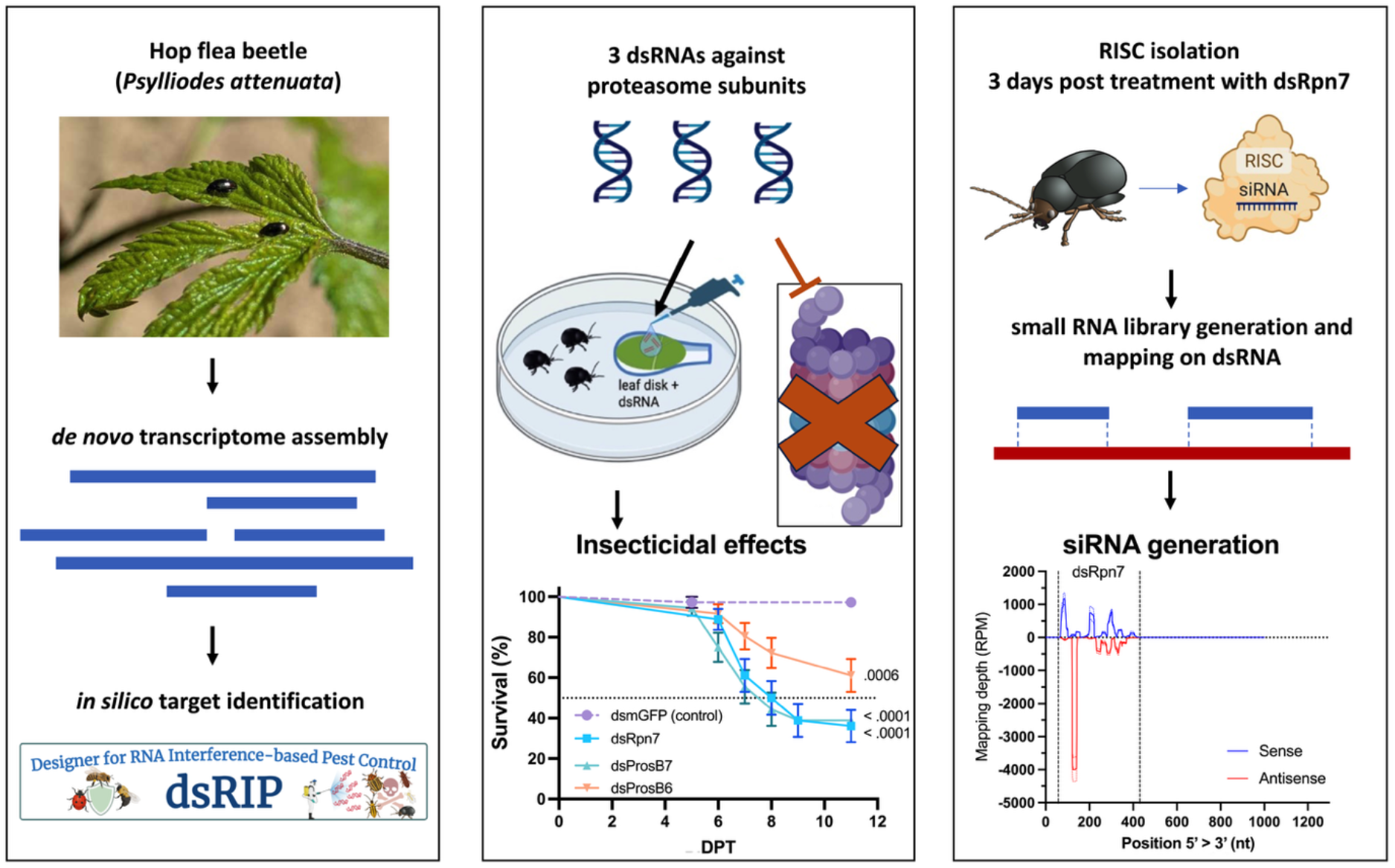

- dsRIP identifies insecticidal proteasome subunit targets in the hop flea beetle.
- Oral dsRNA delivery causes 64% mortality and reduces feeding by up to 45%.
- Ingested dsRNA yields predominantly 21-nt siRNAs associated with RISC.
- A new hop flea beetle transcriptome has 89% BUSCO completeness.

## 1. Introduction

Insect pests cause yield losses of more than 20% in important crops, which are projected to increase due to global warming (Deutsch et al., 2018). Overreliance on chemical insecticides with low species specificity for controlling pest populations has led to an environmental burden and the emergence of insecticide-resistant pest populations (Chen et al., 2023; Ihara and Matsuda, 2018; Lundin, 2021; Willis et al., 2020). To meet the demand for sustainable pest management strategies, biotechnological solutions including RNA interference (RNAi)-based strategies are actively being developed against pests (Baum et al., 2007; Huvenne and Smagghe, 2010; Rodrigues et al., 2021).

RNAi is a conserved pathway in eukaryotes that plays important roles in the defense against viruses and transposons, especially in arthropods (Bonning and Saleh, 2021; Galiana-Arnoux et al., 2006). RNAi is initiated upon the recognition of double-stranded RNA (dsRNA) in the cytoplasm. Perfect or near-perfect dsRNA is processed by the endonuclease Dicer-2 into duplexes approximately 21 nucleotides (nt) long referred to as small interfering RNAs (siRNAs) with characteristic 2-nt 3’ overhangs (Kim et al., 2006; Naganuma et al., 2021). Subsequently, the RNA-induced silencing complex (RISC) is assembled via the incorporation of the guide siRNA strand onto Argonaute-2 (Iwakawa and Tomari, 2021; Kawamata and Tomari, 2010). RISC finds and cleaves RNA molecules complementary to the guide siRNA strand, thereby leading to gene-specific silencing (Iwakawa and Tomari, 2021).

In the context of pest control, dsRNA complementary to essential pest genes is delivered through the oral route, either via foliar dsRNA sprays or transgenic plants transformed with dsRNA-producing constructs (Reinders et al., 2023; Rodrigues et al., 2021). Upon feeding on dsRNA-containing plant tissue, dsRNA is taken up into the cytoplasm primarily via endocytosis and silences essential genes via the RNAi pathway (Zhu and Palli, 2020). The silencing of the essential gene leads to suppression of the corresponding protein, which in turn results in lethal and sublethal effects, including feeding and mobility suppression (Cedden et al., 2024; Mehlhorn et al., 2021). The promise of RNAi-based pest control is exemplified by the development of the foliar dsRNA-based active ingredient Ledprona against the Colorado potato beetle *Leptinotarsa decemlineata* (Rodrigues et al., 2021) and transgenic maize expressing dsRNA complementary to the Snf7 gene of *Diabrotica virgifera virgifera* (Reinders et al., 2023). RNAi-based pest control was also shown to be effective in the cabbage stem flea beetle *Psylliodes chrysocephala* in laboratory bioassays (Cedden et al., 2025c, 2024). A robust trend in RNAi-based pest control is that certain coleopteran families, including the family Chrysomelidae (leaf beetles), respond strongly to orally delivered dsRNA, making them promising targets for this strategy, whereas other orders, such as lepidopterans, exhibit little to no response to exogenous dsRNA (Baum et al., 2007; Terenius et al., 2011; Willow and Veromann, 2021).

Recently, a genome-wide RNAi screen in the red flour beetle *Tribolium castaneum* led to the identification of the most effective (i.e., highly lethal upon knockdown) RNAi target pathways and genes in coleopteran pests (Buer et al., 2025; Cedden and Bucher, 2025). For instance, multiple genes involved in the proteasome pathway, which is responsible for the recycling of dispensable or misfolded proteins, have turned out to be among the most effective targets in *T. castaneum*. Moreover, most proteasome targets identified in *T. castaneum* also turned out to be highly effective targets in multiple coleopteran pests, including *L. decemlineata* and *P. chrysocephala*, indicating a high level of transferability (Cedden et al., 2025a; Cedden and Bucher, 2025). Following the identification of the most effective RNAi targets, recent work has systematically established optimal dsRNA sequence design rules based on the properties of siRNAs processed from the delivered dsRNA in insects (Cedden et al., 2025b). By combining these experimental insights, the dsRIP web platform (https://dsrip.uni-goettingen.de/) offers a complete pipeline for finding the most effective target gene orthologs in pests and designing dsRNA with high predicted efficacy and minimal predicted off-target effects on non-target organisms, such as honey bees.

The hop flea beetle, *Psylliodes attenuata* (Coleoptera: Chrysomelidae), is a univoltine leaf beetle native to the Palaearctic Region, feeding mainly on hops (*Humulus lupulus*) and *Cannabis* (Cannabaceae) (Douglas et al., 2024; Kovaříková et al., 2025). Adults damage hop leaves, lateral shoots, flowers, and cones, while larvae feed below ground on roots. Damage caused by adults may lead to up to 1 week of growth stunting in hop plants. Heavy infestations can reduce cone quantity and quality, causing up to 15% yield losses (Douglas et al., 2024; Kovaříková et al., 2025). Long considered a European and Asian pest, *P. attenuata* has recently been detected in Canada, where populations are now established in Ontario and Québec (Douglas et al., 2024). This expansion highlights the beetle’s capacity to colonize new hop-growing regions and its threat to global hop cultivation, which is crucial for the brewing industry.

In this study, we hypothesized that *P. attenuata* might be susceptible to RNAi-based pest control, given that other beetle species that also belong to Chrysomelidae and the genus Psylliodes exhibit excellent responses to orally delivered dsRNA (Baum et al., 2007; Buer et al., 2025; Cedden et al., 2025a, 2024). To test this idea, we assembled a transcriptome from adult *P. attenuata*, identified effective target genes via the dsRIP web platform, conducted dsRNA feeding bioassays and performed RISC-bound small RNA sequencing to confirm dsRNA processing by the beetle. Our results highlight the potential of proteasome subunits as effective targets for inducing mortality and suppressing feeding, providing a foundation for the development of RNAi-based strategies against this important hop pest.

## 2. Materials and methods

### 2.1. Transcriptome assembly using public data

Initially, public RNA-seq data of adult *P. attenuata* from NCBI BioProject PRJNA768846, SRA SRR16339639 (Gikonyo et al., 2024) were used as input for *de novo* transcriptome assembly using Trinity (v2.15) (Grabherr et al., 2011). Assembly quality was assessed with BUSCO (v5) (Manni et al., 2021) against the endopterygota odb10 dataset. Functional annotation was performed via BlastKOALA (https://www.kegg.jp/blastkoala) (Kanehisa et al., 2016) and the dsRIP web platform (https://dsrip.uni-goettingen.de) (Cedden et al., 2025b).

### 2.2. Orthology inference and target selection

Orthology inference between transcripts from *T. castaneum* and *P. attenuata* was conducted using the pipeline provided by the dsRIP web platform (v1.2, https://dsrip.uni-goettingen.de/). First, the transcripts were converted to proteins based on the longest open reading frame using TransDecoder (v5.7.1, https://github.com/TransDecoder/TransDecoder) and used as input for orthology inference. Candidate essential genes were identified using the dsRIP web platform, with a focus on orthologs of validated RNAi targets in *P. chrysocephala* (Cedden et al., 2025a, 2025c) and highly transferable proteasome subunit targets. Based on these criteria, regulatory particle non-ATPase 7 (*Pa-rpn7*), proteasome beta subunit 6 (*Pa-prosβ6*), and proteasome beta subunit 7 (*Pa-prosβ7*) were selected.

### 2.3. dsRNA design and synthesis

For the bioassays in this study, we used dsRNAs designed by the dsRIP web platform with the following parameters: 300–400 bp length range, restricted to the open reading frame, and 100% priority to efficacy. Target mRNA-specific primers with the T7 promoter added to the 5’ end were designed using the dsRIP web platform and provided by Integrated DNA Technologies (IDT, Germany).

cDNA was obtained from *P. attenuata* adults collected in April by isolating RNA from 5–10 adults using the Quick-RNA Tissue/Insect Kit (Zymo Research, Germany) and the LunaScript® RT SuperMix Kit according to the manufacturers’ instructions. The cDNA was used to amplify the target region with T7 promoter-containing primers via PCR using the Phusion® High-Fidelity PCR Kit (50 µL reaction volume). To obtain the control treatment, dsmGFP, gBlock DNA provided by IDT was used as the template and it was similarly amplified using T7 promoter-containing primers. The PCR annealing temperature was 60 °C and 30 cycles were performed. PCR products were run on a 1% agarose gel and the single bands were excised and purified using the Gel and PCR Clean-up kit (Macherey–Nagel, Germany). The purified PCR products (∼500 ng) were used as templates for in vitro transcription reactions using the MEGAscript™ T7 Transcription Kit (20 µL reaction volume). dsRNAs were purified through lithium chloride precipitation and washed three times with 70% ethanol. Following purification, dsRNAs were reannealed by denaturation at 94 °C for 5 min, followed by cooling at room temperature for 30 min, and were suspended in nuclease-free water. The lengths of the annealed dsRNAs were verified on a 1% agarose gel.

### 2.4. dsRNA feeding bioassays

*P. attenuata* adults were collected in August from experimental hop fields that do not receive insecticide applications (Wolnzach, Germany, 2025) for dsRNA feeding bioassays. Adults were placed in each Petri dish (35 × 10 mm with vents). Next, 3 µL of 500 ng/µL dsRNA solution containing 200 ppm of Triton X-100 (Sigma-Aldrich, Germany) was applied to 37 mm^2^ leaf disks punched from *Humulus lupulus* leaves. Hence, the leaf disks contained dsRNAs at 40.5 ng/mm^2^. The leaf disks were placed on top of 50 mm^3^ 1% agarose gel disks to preserve freshness. After letting the dsRNA solution on the leaf disks dry for 10 min, they were placed inside the Petri dishes together with the adult beetles. After 2 days, the dsRNA-treated leaf disks were replaced daily with fresh untreated leaf disks of the same size until 11 days post treatment. In total, 36 mixed-sex adult beetles were used in the survival bioassays and received either proteasome-targeting dsRNA treatment or a dsmGFP control. Individual beetles were used as data points for the survival analysis. Survival curves of the dsRNA treatment groups were compared with that of the dsmGFP control group using the log-rank test.

To measure the effect of dsRNA treatments on feeding behavior, 10 Petri dishes were prepared as described above. The remaining areas of untreated leaf disks provided on days 2, 3, and 4 were recorded on the subsequent day. The leaves were flattened beside a 5 mm reference line and pictures were taken using a 25-megapixel camera. The remaining leaf area was measured using the ImageJ software (v1.8, https://imagej.nih.gov/ij/), using the ‘analyze particles’ function and subtracted from the initial leaf disk area (37 mm^2^) to obtain consumed leaf area (mm^2^) at 3, 4 and 5 days post treatment (DPT). The consumed leaf area data were analyzed using two-way ANOVA, followed by Dunnett’s test against the control dsmGFP group.

### 2.5. RISC-bound small RNA sequencing

Adult insects were sampled 3 days after ingestion of dsRpn7- or dsmGFP-treated leaf disks. For dsRpn7 treatments, three biological replicates were collected, and for dsmGFP controls, two biological replicates were collected, with each biological replicate containing three P. attenuata adults. Insects were flash-frozen in liquid nitrogen and stored at −80 °C until processing. RISC complexes were isolated using the TraPR Small RNA Isolation Kit according to the manufacturer’s instructions. Frozen samples were homogenized in 300 µL lysis buffer using RNase-free micro pestles and vortexed for 20 s. Lysates were clarified by centrifugation at 10,000 × g for 5 min at +4 °C. Cleared lysates were applied to TraPR columns, mixed with the resin by vigorous inversion for 20 s, and RISC complexes were eluted by centrifugation at 1,000 × g for 15 s. Elution was repeated twice with 250 µL elution buffer per step, yielding 750 µL of RISC-containing eluate. RISC-bound small RNAs were extracted from the eluate by phenol (pH 4.3)/chloroform/isoamyl alcohol precipitation (ROTI®Aqua-P/C/I, Carl Roth GmbH, Germany). Sodium acetate and carrier provided with the kit were used to facilitate small RNA precipitation. Pellets were washed three times with 80% ice-cold ethanol and resuspended in 10 µL RNA elution buffer.

Small RNA libraries were prepared using the NEBNext® Multiplex Small RNA Library Prep Kit (NEB #E7560S) following the manufacturer’s instructions. A total of 6 µL of purified small RNA (∼500 ng RNA) was used as input. Adaptors were used undiluted due to the high small RNA input. Libraries were amplified using 11 PCR cycles, purified with the Monarch PCR & DNA Cleanup Kit (NEB) using a 7:1 binding buffer-to-sample ratio, and eluted in nuclease-free water. Library quality was assessed using an Agilent 2100 Bioanalyzer before and after size selection of 135–150 bp cDNA fragments (corresponding to 16–30 nt inserts) via 6% polyacrylamide gel electrophoresis. Libraries were pooled and sequenced on a DNBSEQ-G400 platform. Raw reads were adapter-trimmed and quality filtered (Phred score >30) using Trim Galore v0.6. Filtered reads were mapped to the corresponding rpn7 or mGFP transcript sequences using Bowtie v1.3, reporting all valid alignments in forward and reverse orientations with no mismatches allowed. Length distributions and read abundances were analyzed from mapped datasets, with particular focus on 21 nt reads.

### 2.6. In silico off-target analysis

Target genes were subjected to *in silico* off-target analysis using the dsRIP web platform. Briefly, this analysis divides the target gene sequence into all possible 21-nt siRNAs, including sense and antisense strands, and searches for matches in the transcriptomes of non-target species. The selected non-target species for this analysis were *Apis mellifera, Bombus terrestris, Coccinella septempunctata, Cotesia congregata, Homo sapiens,* and *Daphnia magna*. Up to 1 mismatch was allowed for a valid match.

### 2.7. RNA sequencing and transcriptome assembly

Total RNA was extracted from 10 field-collected *P. attenuata* adults using the Quick RNA Tissue/Insect Kit (Zymo Research) and purified with the TURBO DNA free kit (Thermo Fisher Scientific). RNA integrity was assessed on an Agilent 2100 Bioanalyzer (Germany) and four biological replicates with RIN values of at least 7.0 were used for library preparation. mRNA libraries were prepared using the NEBNext PolyA mRNA Magnetic Isolation Module followed by the NEBNext kit (E7490). Library quality and quantity were evaluated on the Agilent 2100 Bioanalyzer. Equimolarly pooled libraries were paired-end sequenced by BGI Genomics Tech Solutions on a DNBSEQ-G400 platform. Transcriptome assembly of the resulting RNA-seq data was performed as described for the public RNA-seq data.

## 3. Results

### 3.1. Transcriptome assembly using public data

Identification of effective target genes and design of complementary dsRNAs require a transcriptome of the species of interest. To that end, we *de novo* assembled an adult *P. attenuata* transcriptome using previously deposited public RNA-seq data (Gikonyo et al., 2024). The assembly contained major expected protein families including those involved in genetic information processing and cellular processes (Fig. 1A). The transcriptome assembly had a BUSCO completeness score of 75.7% (single: 58.5% and duplicated: 17.2%; fragmented: 12.9% and missing: 11.4%) (Fig. 1B). Orthology inference via the dsRIP web platform (https://dsrip.uni-goettingen.de) identified *P. attenuata* orthologs of the effective target genes previously identified in *T. castaneum* and successfully transferred to *P. chrysocephala*.

**Figure 1.**
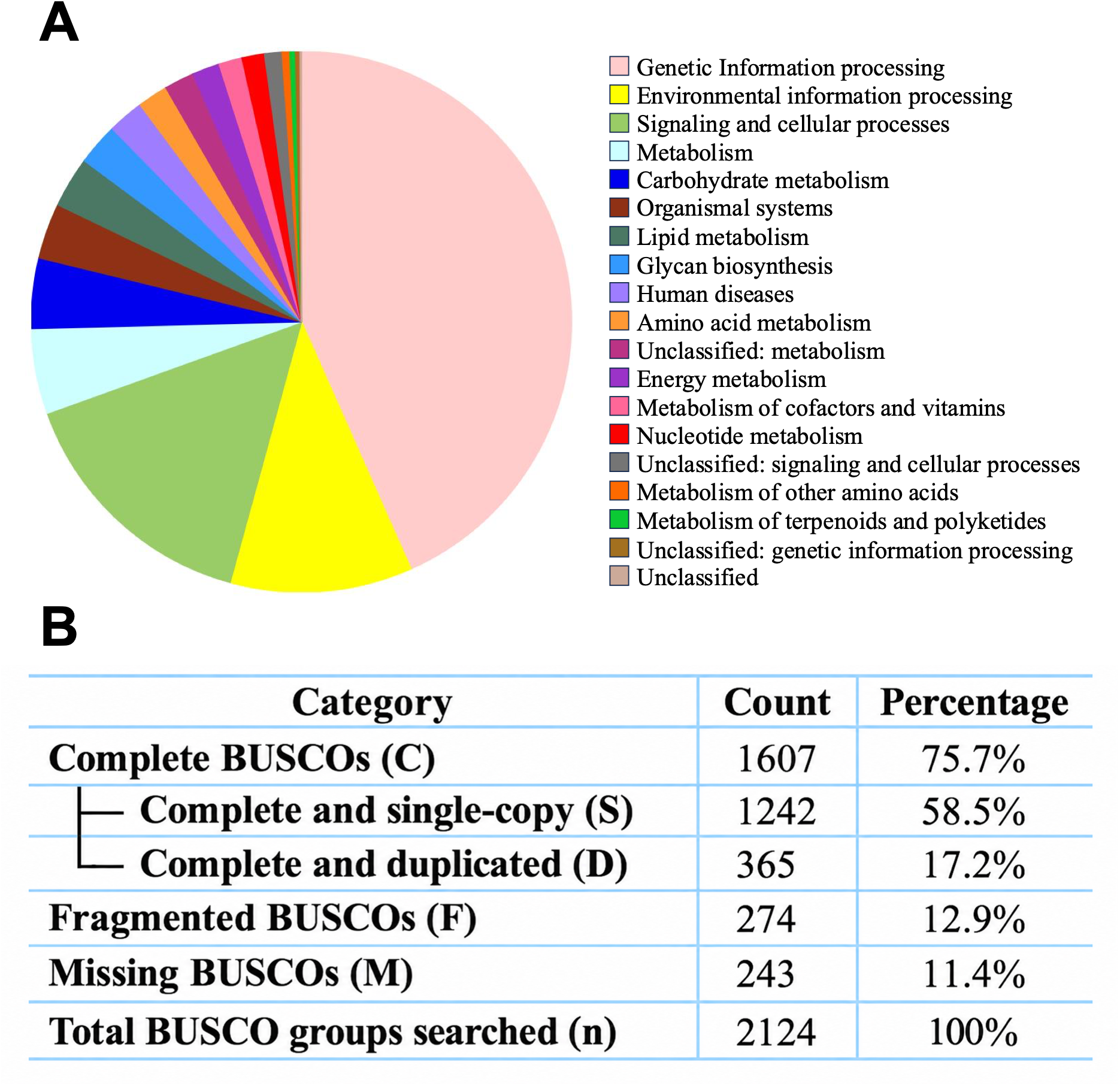
*De novo* transcriptome assembly of *P. attenuata using public RNA-seq*. (A) Distribution of gene families in the de novo transcriptome assembly was investigated using BlastKOALA (https://www.kegg.jp/blastkoala/). (B) BUSCO completeness analysis indicated 75.7% completeness when compared against the endopterygota odb10 dataset.

### 3.2. Candidate gene identification and dsRNA design

Three candidates were selected for experimental testing (Table 1): regulatory particle non-ATPase 7 (*Pa-rpn7*), proteasome beta subunit 7 (*Pa-prosβ7*), and proteasome beta subunit 6 (*Pa-prosβ6*). The selection was based on previous studies in closely related species that identified the proteasome pathway as a highly effective target across coleopteran pests (Buer et al., 2025; Cedden and Bucher, 2025).

**Table 1.**
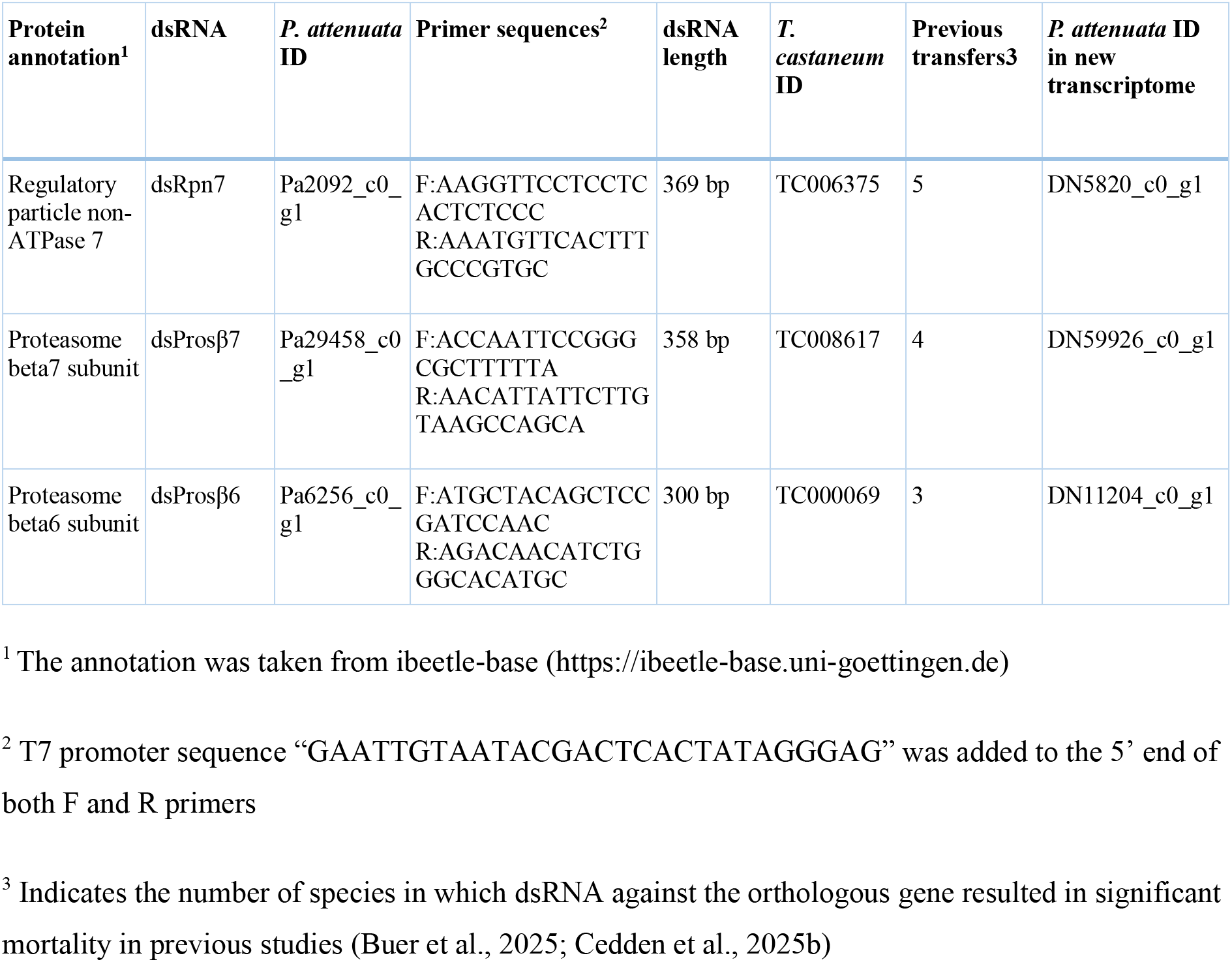
Details of target genes in *P. attenuata* and primer sequences used for dsRNA synthesis.

The next step in the dsRIP pipeline is the design of an effective dsRNA sequence, which is based on the predicted features of all possible siRNAs that could be generated from the dsRNA. We restricted the dsRNA target region to the open reading frame (ORF), one of the efficacy criteria, and selected the region with the highest efficacy score using the default parameters on the dsRIP web platform. The dsRNAs ranged from 300 to 369 bp and included multiple effective siRNA peaks (Fig. 2A–D).

**Figure 2.**
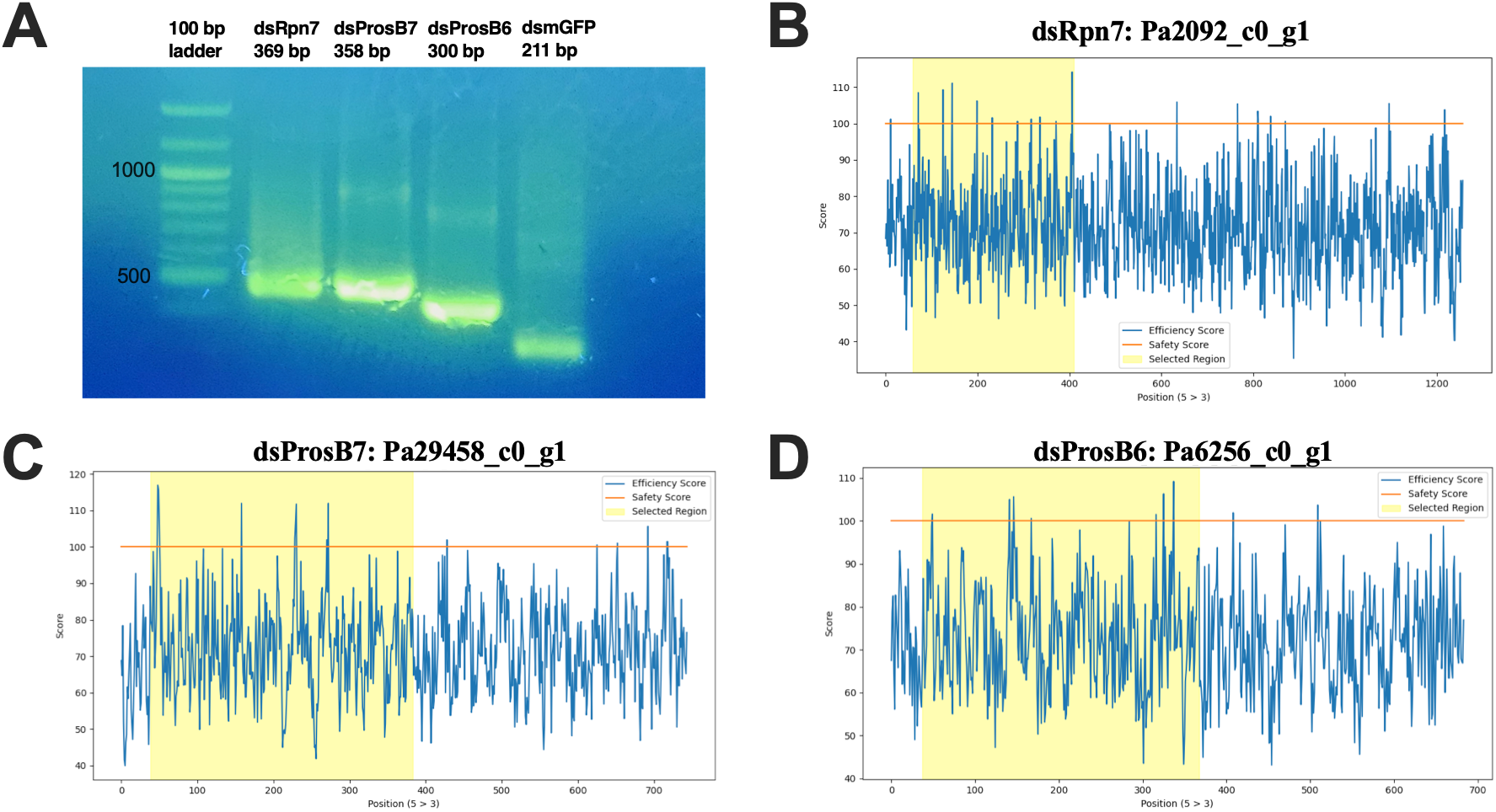
Design and synthesis of *P. attenuata* dsRNAs. (A) Gel electrophoresis of in vitro transcribed dsRNAs targeting Pa-rpn7 (369 bp), Pa-prosβ7 (358 bp), Pa-prosβ6 (300 bp), and dsmGFP control (211 bp). (B–D) Outputs of the dsRIP web platform (https://dsrip.uni-goettingen.de/) instructed to design dsRNA with the highest RNAi efficacy (safety assessment was skipped).

### 3.3. Effects of dsRNAs on adult survival

Next, we conducted dsRNA feeding bioassays to assess effects on survival (Fig. 3A). To that end, adult *P. attenuata* were fed with hop leaf disks treated with dsRNA solution at 40.5 ng/mm^2^ for 2 days and then provided with untreated hop leaf disks. We did not record noticeable effects on survival until 5 DPT for any of the dsRNA treatments, and control adults (dsmGFP) maintained >95% survival throughout the 11-day assay. dsRpn7 and dsProsβ7 had similar, significant negative effects on survival, reaching up to 64% and 61.1% mortality at 11 DPT (P < 0.0001 by log-rank test against the dsmGFP group, n = 36). dsProsβ6 treatment also significantly lowered survival, albeit only by 39% at 11 DPT (P = 0.0006 by log-rank test, n = 36). These results show that orally administered dsRNAs targeting the proteasome components are able to lower the survival of *P. attenuata* adults.

**Figure 3.**
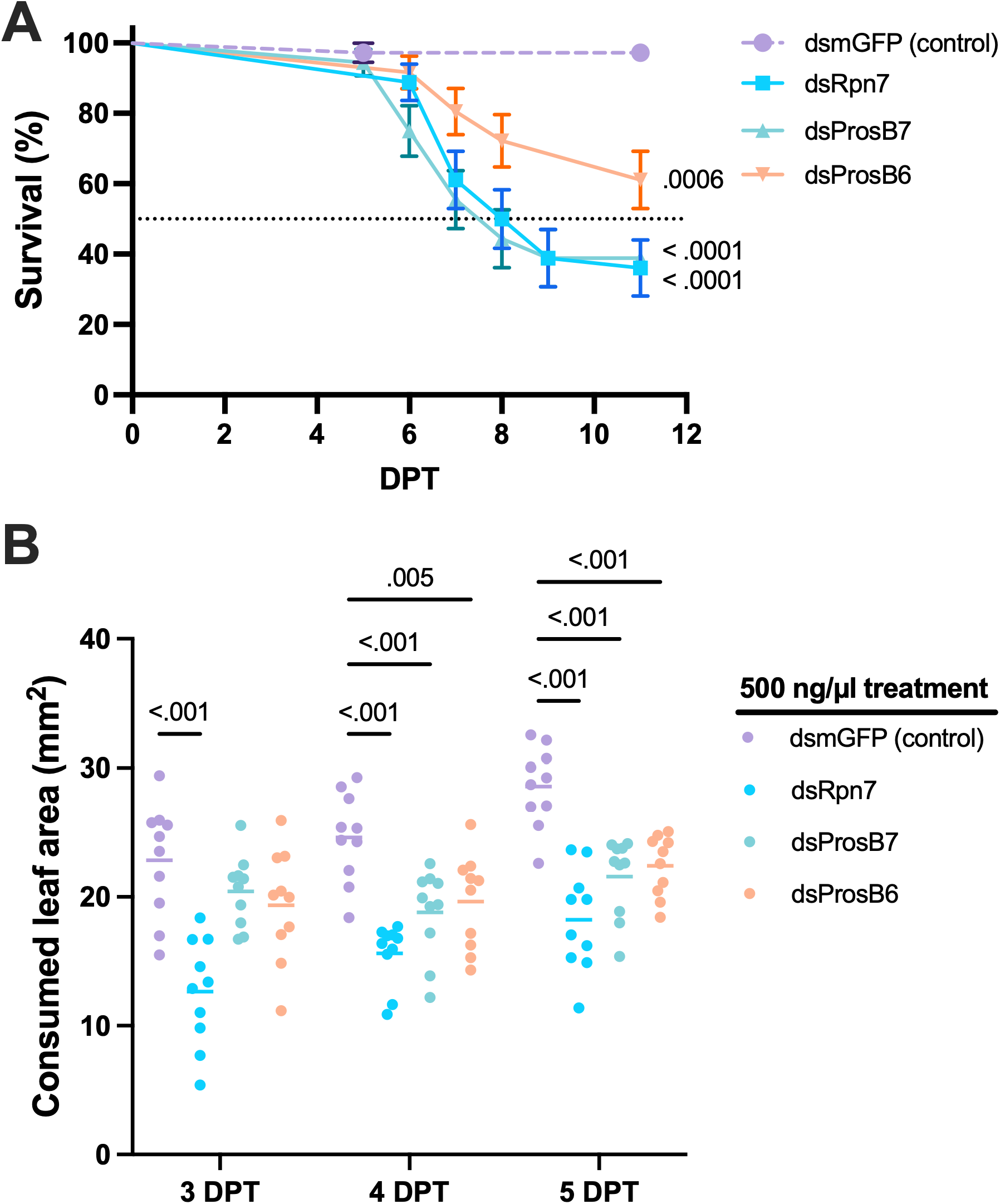
RNAi bioassays in *P. attenuata*. (A) Survival of adult beetles (n = 36, mixed-sex) after oral delivery of 40.5 ng/mm² dsRNA applied on hop leaf disks. P values were calculated by the log-rank test against the dsmGFP control group. (B) Feeding inhibition measured as consumed leaf area in mm² (n = 10 Petri dishes). P values were calculated by Dunnett’s test against the dsmGFP control group. Error bars show the standard error of the mean (SEM) and the horizontal lines indicate the mean.

### 3.4. Effects of dsRNAs on feeding behavior

Insecticidal dsRNA treatments typically result in feeding inhibition that is observable much earlier than effects on survival. Here, we measured the leaf area consumed by the adults from 3 DPT to 5 DPT (Fig. 3B). At 3 DPT, the dsRpn7 group consumed 45% less leaf area compared to the dsmGFP group (P < 0.001 by Dunnett’s test), while the dsProsβ6 and dsProsβ7 groups did not significantly differ from the dsmGFP group (P > 0.05 by Dunnett’s test). At 4 DPT and 5 DPT, all dsRNA treatment groups had lower leaf consumption compared to the dsmGFP group (P ≤ 0.005) with dsRpn7 having the greatest effect on both days (36.5% and 36.2%, respectively). These results show that orally delivered dsRNAs are able to inhibit the feeding behavior of *P. attenuata* prior to the onset of the effect on survival.

### 3.5. RISC-bound small RNA sequencing following dsRNA feeding

The insecticidal effects observed in dsRNA feeding assays prompted us to confirm the activation of the RNAi pathway at the molecular level. To that end, we conducted RISC-bound small RNA sequencing 3 days following dsRpn7 or dsmGFP feeding to identify dsRNA-derived siRNAs. The results showed that dsRpn7 was processed predominantly into ∼21 nt siRNAs mapping onto the dsRNA target region, but not outside the target region in the rpn7 transcript (n = 3, Fig. 4A). We observed a major antisense (i.e., complementary to the target transcript) siRNA hotspot around 120 nt of the *rpn7* transcript, which is likely to have mainly contributed to the insecticidal effects of the dsRNA. Antisense siRNA peaks were expected as the dsRNA was optimized using the dsRIP web platform to generate effective siRNAs. Similarly, the negative control dsmGFP was also processed into dsmGFP-mapping ∼21 nt siRNAs (n = 2, Fig. 4B). The finding that both dsRNAs are processed mainly into 21 nt siRNAs (Fig. 4C and 4D) is in accordance with studies in other coleopteran pests where injected or fed dsRNA is also processed primarily into 21 nt siRNAs. The lack of siRNA length peaks above 24 nt suggests that the delivered dsRNA was not processed by the Piwi-interacting RNA pathway.

**Figure 4.**
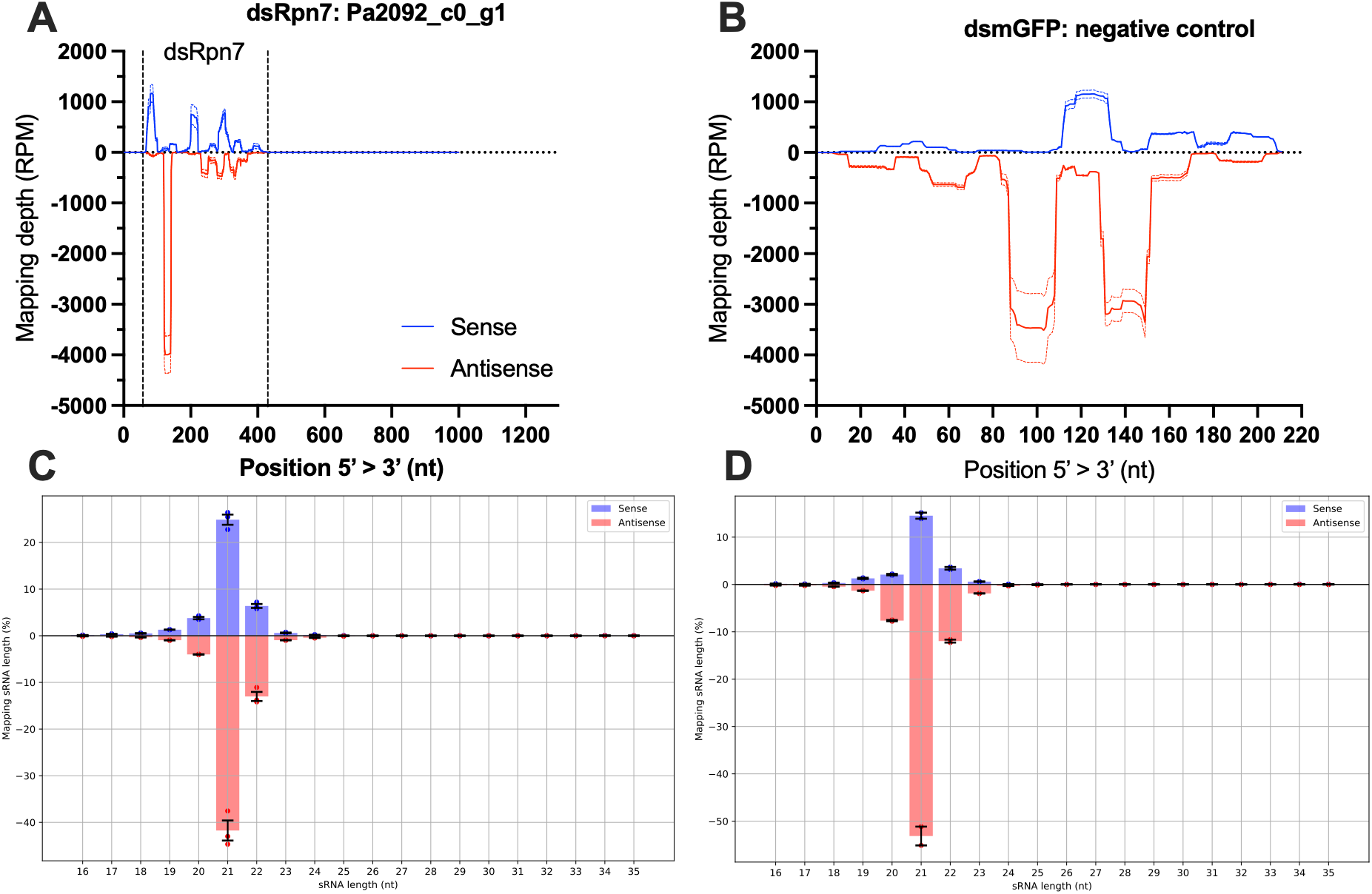
RISC-bound small RNA sequencing following dsRNA feeding. (A) Distribution of RISC-associated small RNAs mapping to the *rpn7* transcript 3 days post dsRpn7 feeding (n = 3). Reads map predominantly within the dsRNA target region, with sense reads shown in blue and antisense reads in red. A prominent antisense siRNA hotspot is observed around position ∼120 nt. (B) Distribution of RISC-associated small RNAs mapping to the mGFP transcript 3 days post dsmGFP feeding (n = 2), showing efficient processing of the control dsRNA into siRNAs. (C–D) Length profiles of RISC-bound small RNAs derived from dsRpn7 (C) and dsmGFP (D), demonstrating a strong enrichment of ∼21 nt siRNAs.

### 3.6. In silico off-target analysis

The bioassays showed that all three proteasome-targeting dsRNAs significantly reduced survival and leaf consumption. Hence, we analyzed the safety profiles of the sequences of these target genes using the dsRIP web platform to assess their suitability for further development (Fig. 5). dsRIP was instructed to assess all possible 21 nt siRNAs that could be generated from the target genes and report potential off-targeting events with up to 1 mismatch in the transcriptomes of important non-target organisms, including pollinators, predators, parasitoids, aquatic invertebrates, and humans. Although *Pa-rpn7* had up to 40 predicted off-target genes for certain siRNA regions, it had no essential gene off-targets in the investigated non-target organisms. In contrast, *Pa-prosβ7* and *Pa-prosβ6* had fewer off-target genes, only up to 5 per siRNA region. However, both had predicted essential off-target genes, with 5 possible siRNAs from *Pa-prosβ6* targeting an *A. mellifera* essential gene ortholog (XM_395163.6). These results highlight the importance of integrating gene essentiality into *in silico* predictions of potential off-target effects on non-target organisms, as this approach prioritizes potential off-targets most likely to pose ecological risk.

**Figure 5.**
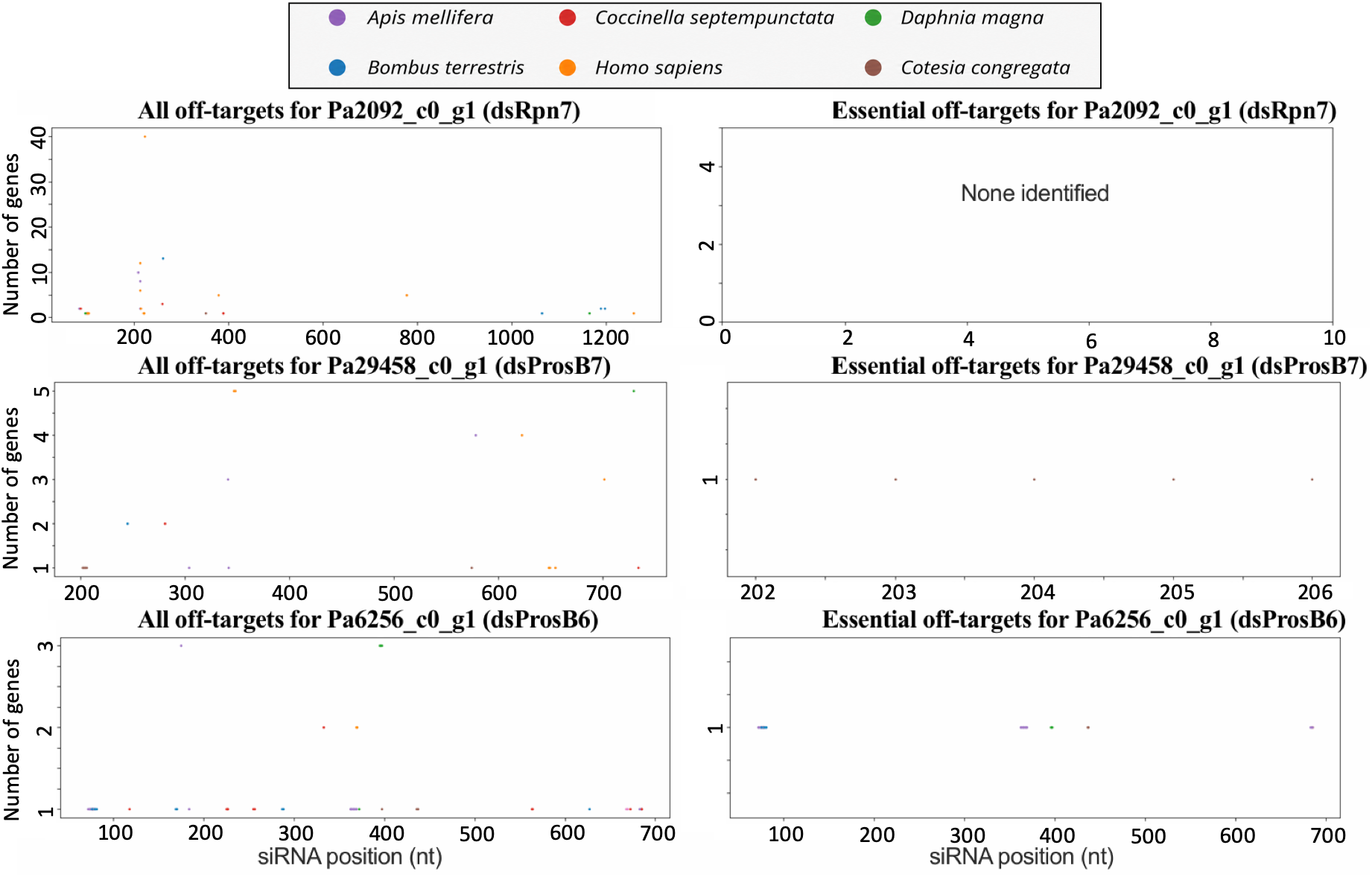
*In silico* safety assessment of effective target genes identified in *P. attenuata*. The safety assessment was conducted using the dsRIP web platform (https://dsrip.uni-goettingen.de). The assessment involves generating all possible 21 nt siRNAs from a gene sequence and analyzing whether they have potential off-targets with up to 1 mismatch in the transcriptomes of selected non-target organisms. Left plots show all predicted off-target genes per siRNA position, while right plots show only those that are orthologous to essential genes identified in *T. castaneum* through a genome-wide RNAi screen.

### 3.7. Higher-quality transcriptome assembly

The dsRNA designs used in this study were based on a transcriptome assembly generated from a single publicly available biological replicate. To assemble a higher-quality transcriptome for future RNAi studies in *P. attenuata*, we generated RNA-seq data from four biological replicates of intact *P. attenuata* adults and used this dataset as input for a new de novo transcriptome assembly. Using the same in silico procedure applied to the public dataset, we assembled 82,242 transcripts with a BUSCO completeness score of 89.0% against the Endopterygota odb10 dataset (Table 2). The improvement from 75.7% in the public data-based assembly to 89.0% completeness suggests that this updated transcriptome may be more useful for future RNAi studies in *P. attenuata*. The newly generated transcriptome also contained the three proteasome subunit targets tested in this study, with 100% sequence similarity for the dsRNA target regions (Table 1).

**Table 2.** BUSCO score of transcriptome assembly of *P. attenuata* adults using new RNA-seq data.

| Category | Count | Percentage |
| --- | --- | --- |
| **Complete BUSCOs (C)** | **1890** | **89.0%** |
| └─Complete and single-copy BUSCOs (S) | 1822 | 85.8% |
| └─Complete and duplicated BUSCOs (D) | 68 | 3.2% |
| **Fragmented BUSCOs (F)** | **99** | **4.7%** |
| **Missing BUSCOs (M)** | **135** | **6.4%** |
| **Total BUSCO groups searched** | **2124** | **100%** |

## 4. Discussion

This study establishes the hop flea beetle *P. attenuata* as a viable target of RNAi-based pest control and identifies three proteasome subunit genes as highly effective for inducing insecticidal effects. The results mirror findings in *P. chrysocephala* and *L. decemlineata*, where proteasome subunits proved the most potent targets (Buer et al., 2025; Cedden et al., 2025a, 2025b; Rodrigues et al., 2021). Such cross-species convergence underscores the potential of RNAi-mediated proteasome inhibition in the control of chrysomelid beetles. Moreover, the study demonstrates the usefulness of the dsRIP web platform in establishing novel pest species for RNAi-based pest control and might act as a guide for efficiently establishing other pests for this purpose.

Here, we showed that up to 64% mortality over 11 days and 45% feeding inhibition as early as 3 DPT can be achieved through a single exposure to dsRpn7-treated leaf disks in *P. attenuata.* Future studies should focus on improving these insecticidal effects through the identification of more effective target genes. Although it has been demonstrated that there is ∼80% transferability of target genes between chrysomelid species (leaf beetles), predicting which target genes will yield the best survival curves, dose-response curves and sublethal effects is challenging (Cedden and Bucher, 2025). This unpredictability stems from the complex factors that determine the phenotype following RNAi of a particular gene. For instance, even for orthologous genes, the mRNA and the protein will vary in their cellular half-lives (Rahman and Sadygov, 2017), with longer half-lives delaying the phenotype following RNAi.

Furthermore, in each species, the gene regulatory network that includes the target gene might slightly differ, which in turn determines whether the target gene expression is more or less susceptible to perturbations (MacNeil and Walhout, 2011). Another likely contributing factor is the varying level of compensatory action by paralogous or other proteins that mitigate the effect of RNAi depending on the species (Cedden and Bucher, 2025). Hence, superior targets for higher insecticidal effects will need to be identified through further experimental screens in *P. attenuata* and previous large-scale RNAi studies in *T. castaneum* and *P. chrysocephala* would be excellent resources, as they were in the present study (Buer et al., 2025; Cedden et al., 2025a). The higher-quality transcriptome generated in this study will facilitate identification of RNAi target genes with higher insecticidal efficacy.

The RISC-bound small RNA-seq experiments further confirmed the promise of RNAi-based pest control against *P. attenuata*, as the processing of orally ingested dsRNA by the RNAi machinery underlies this strategy. The similar patterns of dsRNA processing in *P. attenuata* compared to other coleopteran pests, including *T. castaneum* and *P. chrysocephala*, suggest that previous approaches for optimizing dsRNA designs against these pests could be readily applied to *P. attenuata*.

The high economic value of hop crops, the recent spread of *P. attenuata* into North America and restrictions on the application of neonicotinoids in Europe make research into its RNAi-based control timely. This work opens up the possibility of developing foliar dsRNA sprays for the management of *P. attenuata*, whose population is expanding into new regions (Douglas et al., 2024). Short-term research goals in this regard include further molecular confirmation of RNAi effects, including investigation of target gene knockdown, identification of more effective target genes and investigation of dose responses to dsRNAs. Long-term research goals include assessing potential effects of dsRNAs on non-target organisms, establishing suitable dsRNA formulations and application strategies on hop plants, and testing such strategies in field trials, where effects of environmental factors, including UV and microbial degradation, on dsRNA application efficacy could be evaluated (Bachman et al., 2020). In particular, the perennial life cycle and height of up to 10 meters of hop plants call for specialized approaches for feasible RNAi-based control of *P. attenuata*.

## 5. Conclusions

This study establishes the hop flea beetle *Psylliodes attenuata* as a promising target for RNAi-based pest control and confirms the proteasome subunits as effective molecular targets for inducing both lethal and feeding-inhibitory effects. By generating a transcriptome and utilizing the recently developed dsRIP web platform, we identified and designed effective dsRNAs that reduced survival and feeding, while *in silico* analyses characterized the safety profiles of the target genes against non-target organisms, such as honey bees. Importantly, this work directly tested the complete dsRIP pipeline in a new pest species with respect to RNAi research, demonstrating its usefulness for rapidly extending RNAi-based strategies to additional insect pests. These results not only support the future development of dsRNA-based sprays or transgenic hop varieties but also may act as a transferable framework for effectively establishing novel pests for RNAi-based pest control.

## CRediT authorship contribution statement

Doga Cedden: Conceptualization, Methodology, Funding acquisition, Investigation, Data curation, Formal analysis, Writing – original draft, Writing – review and editing. Gözde Güney: Investigation, Writing – review and editing. Paul Plathner: Data curation, Formal analysis, Writing – review and editing. Michael Rostás: Resources, Writing – review and editing. Stefan Scholten: Resources, Writing – review and editing. Christina Krönauer: Conceptualization, Methodology, Funding acquisition, Writing – review and editing.

## Data availability

Raw data supporting the findings of this study are provided in Supporting File 1 and are also available on Zenodo (https://doi.org/10.5281/zenodo.23457913). De novo transcriptome assemblies are provided as Supporting Files 2 and 3. Raw RNA-seq (BioProject: PRJNA1366906) and small RNA-seq (BioProject: PRJNA1400957) files were deposited in the NCBI SRA database.

## Acknowledgements

The authors thank Prof. Dr. Gregor Bucher for feedback on the manuscript.

